# Development of follicular dendritic cells in lymph nodes depends on retinoic acid mediated signaling

**DOI:** 10.1101/2020.05.20.106385

**Authors:** Jasper J. Koning, Anusha Rajaraman, Rogier M. Reijmers, Tanja Konijn, Junliang Pan, Carl F. Ware, Eugene C. Butcher, Reina E. Mebius

**Affiliations:** Department of Molecular Cell Biology and Immunology, Amsterdam UMC, Vrije Universiteit Amsterdam, Amsterdam Infection and Immunity Institute, Amsterdam, the Netherlands; Laboratory of Immunology and Vascular Biology, Department of Pathology, Stanford University School of Medicine, Stanford, California, USA; Palo Alto Veterans Institute for Research, Palo Alto, California, USA; Sanford Burnham Medical Research Institute, Infectious and Inflammatory Diseases Research Center, Laboratory of Molecular Immunology, La Jolla, USA; The Center for Molecular Biology and Medicine, Veterans Affairs Palo Alto Health Care System, Palo Alto, California, USA

## Abstract

Specialized stromal cells occupy and help define B- and T cell domains, which is crucial for proper functioning of our immune system. Signaling through lymphotoxin and TNF-receptors is crucial for development of different stromal subsets which are thought to arise from a common precursor. However, mechanisms that control the selective generation of the different stromal phenotypes are not known.

Here we show that in mice, retinoic acid mediated signaling is important for the differentiation of precursors towards the Cxcl13^pos^ follicular dendritic cell (FDC) lineage, while blocking lymphotoxin mediated Ccl19^pos^ fibroblastic reticular cell (FRC) lineage differentiation. Consequently, we see at day of birth Cxcl13^pos^Ccl19^neg/low^ and Cxcl13^neg/low^Ccl19^pos^ cells within neonatal lymph nodes.

Furthermore, ablation of retinoic acid receptor signaling in stromal precursors early after birth reduces Cxcl13 expression, while in addition, complete blockade of retinoic acid signaling prevents formation of FDC networks in lymph nodes.

## Introduction

Lymph nodes are strategically located to enable quick and effective interactions between antigen presenting cells and lymphocytes leading to antigen specific immune responses.

The non-hematopoietic stromal cells are important contributors to lymph node homeostasis as well as immune responses (Chang & Turley, 2015; Krishnamurty & Turley, 2020). Stromal cells provide an architecture with distinct microdomains for both B and T lymphocytes that mediate survival(Link et al., 2007; Mueller & Germain, 2009; Takada & Jameson, 2009), tolerance to self-antigens (A.P. Baptista et al., 2014; Fletcher et al., 2010; Lee et al., 2007; Magnusson et al., 2008; Nichols et al., 2007), while in addition they can crucially influence ongoing immune responses (Ahrendt, Hammerschmidt, Pabst, Pabst, & Bode, 2008; A. P. Baptista et al., 2019; Hammerschmidt et al., 2008; Molenaar et al., 2009; Wolvers et al., 1999).

The stromal cell compartment can be subdivided into various endothelial and mesenchymal subsets (Mueller & Germain, 2009). Until recently, the major different mesenchymal stromal subsets included fibroblastic reticular cells (FRCs), marginal reticular cells (MRCs) and follicular dendritic cells (FDCs) (Chang & Turley, 2015; Koning & Mebius, 2012). Additional mesenchymal stromal subtypes were identified using single cell sequencing (Rodda et al., 2018) and have shed light on the transcriptional profile of lymph node stromal cells during homeostasis and upon immune activation. Despite this increased knowledge, it remains unknown which signaling events drive the differentiation from precursor cells into these various stromal clusters.

From previous studies we know that precursors for stromal cells, including FDCs, are already present in mesenteric lymph nodes of newborn mice (Cupedo, Jansen, Kraal, & Mebius, 2004). Recently, we have shown that nestin^pos^ precursors give rise to the various endothelial and mesenchymal derived lymph node stromal cells (Koning et al., 2016).

More specific studies on the origin of FDCs provided evidence for the presence of local mesenchymal FDC-precursors both in spleen and lymph nodes (Castagnaro et al., 2013; Jarjour et al., 2014; Krautler et al., 2012). The presence of B cells is essential for the development of FDCs and has been shown to rely on lymphotoxin and TNF receptor signaling (Endres et al., 1999; Fu, Huang, Wang, & Chaplin, 1998). The presence of perivascular precursors for FDCs in the spleen appeared to be independent of lymphotoxin signaling (Krautler et al., 2012) suggesting that a sequence of signaling events is needed for their differentiation and final maturation to become FDCs.

Here we provide evidence that post-natal retinoic acid signaling critically controls development of FDCs. We identify endothelial cells as additional cellular source for the production of retinoic acid thereby highlighting the importance of endothelial and non-endothelial stromal cells for proper development of lymph node niches. Cell specific inhibition of retinoic acid signaling in nestin^pos^ precursor cells reduced Cxcl13 expression but not FDC development. However, upon pharmacological inhibition of retinoic acid signaling we could show that the appearance of FDCs critically depends on retinoic acid signaling during neonatal lymph node development.

## Results

### Differential stimulation via receptors for retinoic acid and lymphotoxin induces distinct gene expression profiles in mesenchymal cells in vitro

Lymph node development during embryogenesis involves direct crosstalk between lymphoid tissue inducer (LTi) cells and different types of stromal organizer cells (Bovay et al., 2018; Mebius, 2003; Onder et al., 2017) and requires signaling mediated by retinoic acid as well as lymphotoxin (Honda et al., 2001; S. A. van de Pavert et al., 2009). Mesenchymal cells at the presumptive lymph node site are initially exposed to retinoic acid after which LTα1β2 expressing LTi cells will trigger LTβR-mediated signaling in these cells, allowing their maturation towards stromal organizer cells (S. A. van de Pavert et al., 2009). To determine whether the activity of these signaling pathways either in sequence or on their own could contribute to the different fates of stromal cell subsets present in adult lymph nodes we set up *in vitro* cultures of early passage E13.5 embryonic mesenchymal cells enriched for mesenchymal stem cells (E13.5 MSC) as confirmed by their tri-lineage differentiation capacity (fig. 1A).

**Figure 1.**
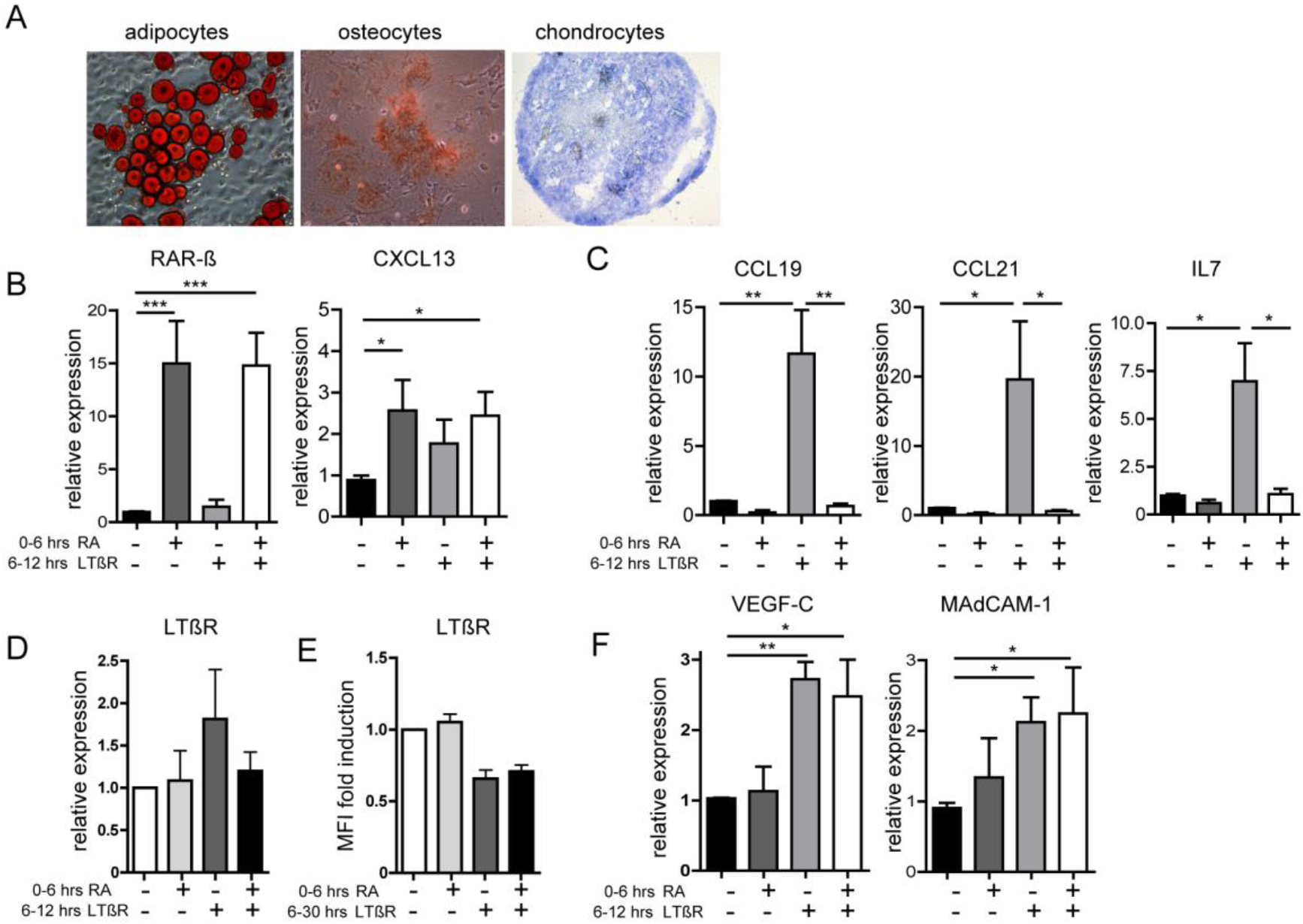
Retinoic Acid signaling prevents upregulation of FRC related genes. A) E13.5 mesenchymal stromal cells (MSC) have the capacity to differentiate into adipocytes, osteocytes and chondrocytes (n=3). B-D, F) mRNA expression of E13.5 MSC upon sequential stimulation with retinoic acid (RA) (6hrs) followed by stimulation with LTβR (6hrs) showing expression of RARβ and Cxcl13 (B, n=9), Ccl19, Ccl21 and Il-7 (C, n=3), LTβR (D, n=3) and VEGF-C and MAdCAM-1 (F, n=3). E) Flow cytometric analysis of LTβR cell surface expression upon stimulation with retinoic acid (6hrs) followed by stimulation with LTβR (30hrs) depicted as fold induction of MFI relative to control. The data represent mean ± SEM; *, p < 0.05; **, p < 0.01; ***, p < 0.001, one-way ANOVA with Bonferroni’s multiple comparison test.

Transcript analysis upon stimulation with retinoic acid alone showed strong upregulation of the β isoform of the receptor (RARβ), which is specifically involved in the induction of Cxcl13 during the initiation phase of lymph node development (S. A. van de Pavert et al., 2009). Indeed, stimulation with retinoic acid induced Cxcl13 expression in mesenchymal precursors while LTβR stimulation alone did not lead to significant upregulation of Cxcl13, although low expression levels could be detected, as shown before (Dejardin et al., 2002) (fig. 1B).

LTβR stimulation (Crowe et al., 1994) initially leads to the activation of the classical Nf-κB pathway (Dejardin et al., 2002; Siebenlist, Franzoso, & Brown, 1994) while upon prolonged triggering, also the alternative Nf-κB pathway becomes activated resulting in the upregulation of Ccl19, Ccl21 and Il-7 via nuclear localization of relB:p52 dimers (Dejardin et al., 2002; Vondenhoff et al., 2009). As expected, LTβR stimulation of mesenchymal precursors resulted in a significant upregulation of Ccl19, Ccl21 and Il-7 transcripts, which are all expressed by FRC in the T cell zone (fig. 1C). Strikingly, cells that were first stimulated with retinoic acid and subsequently via LTβR completely failed to induce Ccl19, Ccl21 and Il-7 transcripts (fig. 1C) but not Cxcl13 (fig. 1B). This inhibition was not the result of downregulation of the receptor itself, since retinoic acid stimulation changed neither mRNA expression nor cell surface expression of LTβR (fig. 1D and E). We also observed no difference in the activation of the non-canonical Nf-kB pathway, since there was similar nuclear relB translocation, while the DNA binding capacity of nuclear relB and p52 was also not affected (suppl. fig. 1). We observed that other LTβR mediated molecules such as VEGF-C and MAdCAM-1 (Vondenhoff et al., 2009) were not affected upon retinoic acid pre-incubation (fig. 1F).

Owing to these observations, we hypothesized that retinoic acid stimulation serves as a fate determining signal that allows the differentiation of stromal precursor cells towards a follicular dendritic cell signature by preventing their differentiation into T cell zone FRCs upon subsequent LTβR triggering.

### Localization of pre-FDCs within post-natal developing lymph nodes

Retinoic acid and LTβR mediated signaling takes place before birth during lymph node development. Therefore stromal cells that have either received initially only retinoic acid mediated signaling, versus stromal cells that received their first signals via LTβR are potentially already present within lymph nodes at day of birth. To address this we monitored mRNA expression of Cxcl13, induced by retinoic acid mediated signaling, as well as Ccl19, expressed in the majority of stromal precursors (Chai et al., 2013) and induced upon LTβR signaling. Upon analysis of wk0 lymph nodes, we observed expression of Ccl19 mRNA as well as Cxcl13 mRNA within cells throughout the lymph node. Although the majority of cells expressed both molecules, cells with high expression of Cxcl13 mRNA appeared to have low amounts of Ccl19 mRNA, and vice versa (fig. 2A).

**Figure 2.**
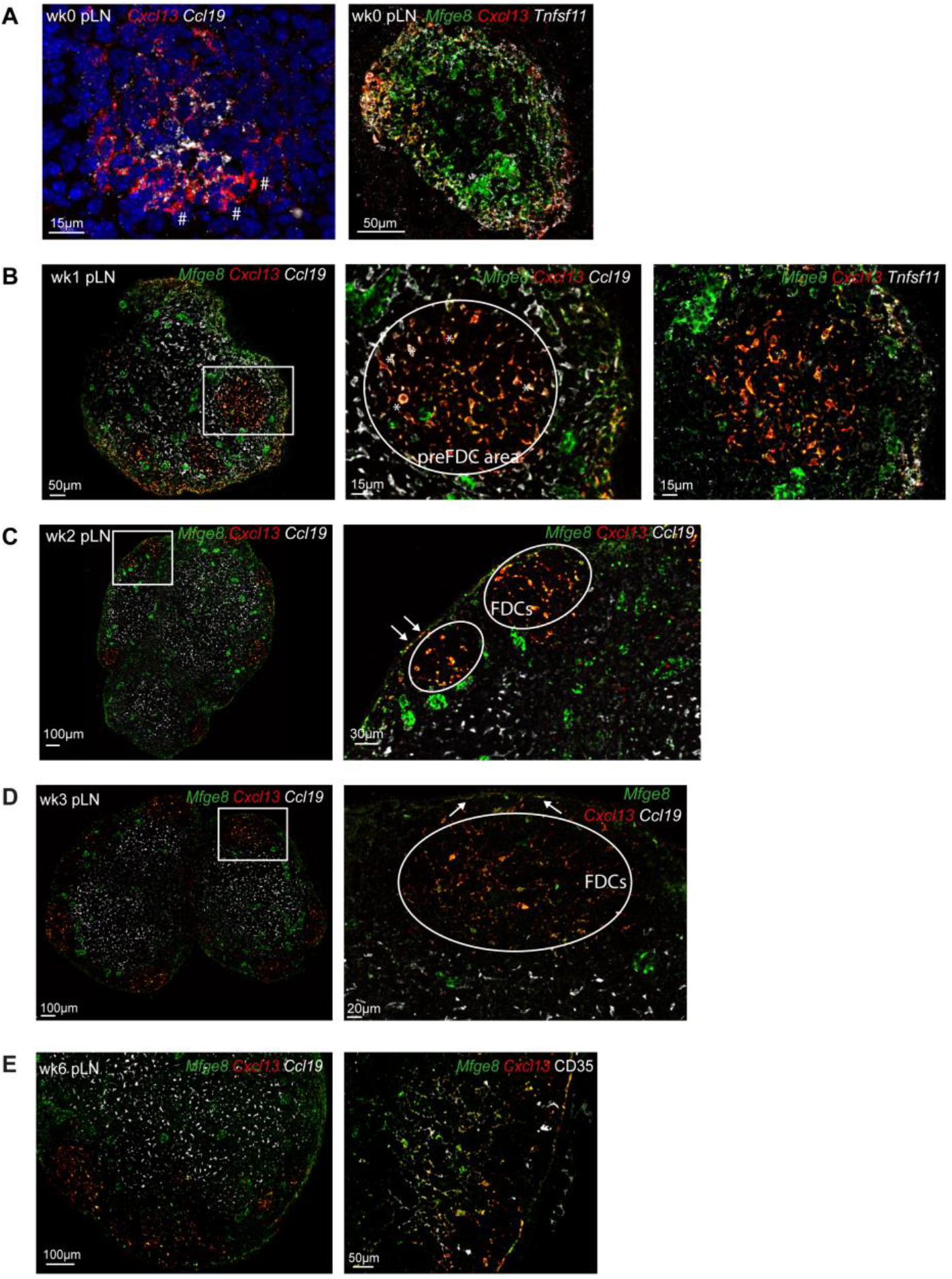
Localization of FDC precursors. A-E) Confocal analysis of multiple fluorescence *in situ* hybridizations at different timepoints after birth in peripheral lymph nodes (pLN). A) Combined expression of Cxcl13 and Ccl19 (left) or Mfge8, Cxcl13 and Tnfsf11 (right) in peripheral lymph nodes at day of birth. Hashtags indicate cells expressing high levels of Cxcl13 and low levels of Ccl19. B) Combined expression of Mfge8, Cxcl13 and Ccl19 (left and middle image) and Mfge8, Cxcl13 and Tnfsf11 (right image) in lymph nodes at 1 week after birth. Asterix indicate Mfge8^pos^Cxcl13^pos^Ccl19^pos^ cells in the preFDC area. C & D) Expression of Mfge8, Cxcl13 and Ccl19 in lymph nodes at week 2 (C) and week 3 (D) after birth. Arrows indicate marginal reticular cells (MRCs). E) Mfge8, Cxcl13 and Ccl19 expression (left) and Mfge8, Cxcl13 and CD35 expression (right) in lymph nodes at 6 weeks after birth. Data are representative for at least n=3 individual mice per timepoint.

We additionally monitored the expression of Mfge8 mRNA, expressed by FDC precursors within the spleen, and Trance (Tnfsf11), expressed by lymphoid tissue organizer (LTO) cells as well as MRCs (Jarjour et al., 2014; Katakai et al., 2004). We observed that all Cxcl13^pos^ cells additionally expressed Mfge8. The strongest levels of Cxcl13 mRNA could be detected in cells that located near the outer border of the lymph node and the majority of the cells expressed Trance mRNA as well defining them as LTO cells with MRC characteristics (fig. 2A). At 1 week after birth, when the first clustering of B cells within lymph nodes has occurred, Cxcl13^pos^Mfge8^pos^ cells were located within follicle areas, as well as in a large area at the outer zone of the lymph node (fig 2B). When combined with Ccl19, some Cxcl13^pos^ cells also expressed Ccl19 within the B cell follicles although the majority was negative for Ccl19. Cxcl13^neg^ cells were mostly excluded from the follicle areas and present within the developing T cell zone where they express Ccl19. The majority of cells within the presumptive B cell follicle did not express Trance mRNA, indicating that they are not MRCs (fig 2B). Within presumptive B cell follicle areas in week 1 lymph nodes, we occasionally observed perivascular located cells that expressed Mfge8, Cxcl13, and Ccl19 (suppl. fig. 2A). Within readily defined B cell follicles at week 2, we observed perivascular cells that were either Mfge8^pos^Cxcl13^pos^ or Mfge8^pos^Ccl19^pos^ cells (suppl. fig. 2B). The separation between Cxcl13^pos^Ccl19^neg^ and Cxcl13^neg^Ccl19^pos^ cells within distinct B-cell areas versus T-cell areas became more prominent when lymph nodes matured further from 2-6 weeks after birth (fig 2C-E).

From this data we can conclude that the cells that received retinoic acid mediated signaling as initial trigger during lymph node development and therefore expressed Cxcl13 but not or low Ccl19, were indeed present within neonatal lymph nodes. With the combined expression of Mfge8 and Trance, these cells have the phenotypic profile of MRCs and are already present at day of birth. Furthermore, in the developing lymph nodes at 1 week of age both Ccl19^pos^ cells, triggered via LTβR, as well as Cxcl13^pos^ cells, triggered via retinoic acid, were located within the developing B cell follicles. The Ccl19^pos^ cells specifically disappeared from the developing B cell follicles in the weeks thereafter.

### Multiple cellular sources of retinoic acid within lymph nodes

To determine whether retinoic acid mediated signaling still occurs after birth and thereby could further contribute to the developing FDC network, we addressed whether cellular sources of retinoic acid were present within neonatal lymph node. Retinoic acid synthesis depends on the 2-step conversion of vitamin A into retinaldehyde and retinoic acid respectively. The final and irreversible step of this conversion involves members of the aldehyde dehydrogenase enzyme family; Aldh1a1, Aldh1a2, Aldh1a3 and expression of these enzymes identifies cells that produce retinoic acid (Kedishvili, 2016). It is known that stromal cells as well as dendritic cells in adult lymph nodes express Aldh1 enzymes (Hammerschmidt et al., 2008; Molenaar et al., 2011), but the expression during early post-natal lymph node development has not been studied before.

Therefore, we sorted the three main stromal subsets, FRCs, BECs and LECs as well as dendritic cells (CD11c^high^MHCII^high^) from pLN of week 1 and week 2 old mice, to examine the expression of Aldh1 enzymes. Dendritic cells are known to express high levels of Aldh1a2 (Zhang et al., 2016) and transcriptomic analysis indeed showed that dendritic cells expressed the highest level of Aldh1a2 at both timepoints in pLN but lacked both Aldh1a1 and Aldh1a3 expression. Sorted FRCs expressed both Aldh1a1 and Aldh1a3 but not Aldh1a2. The expression of Aldh1a1 was higher in week 2 sorted FRCs. To our surprise, we found that BECs also expressed Aldh1a1, while LECs express Aldh1a3 (fig. 3A and supplementary fig 3).

**Figure 3.**
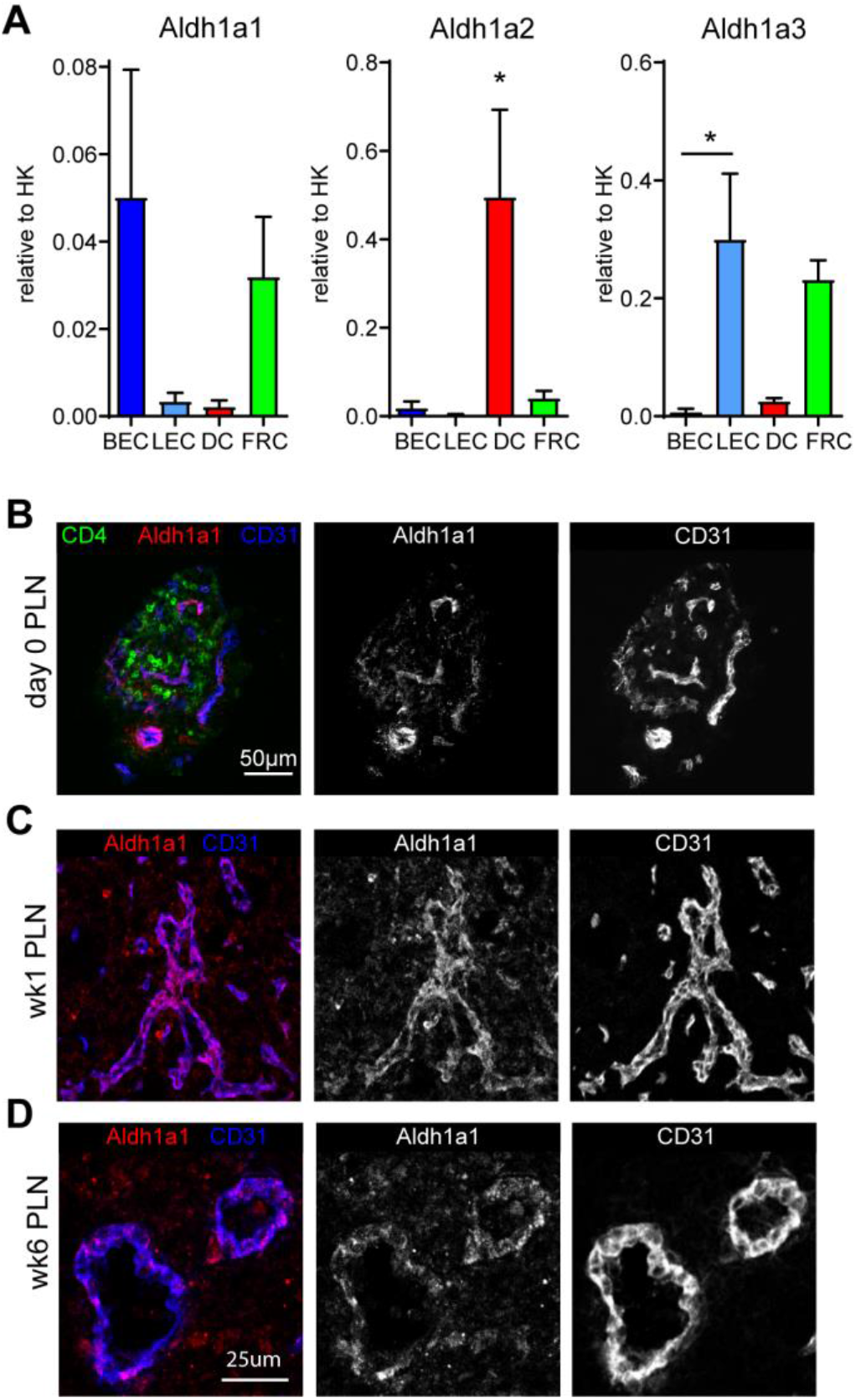
Multiple cellular sources of Aldh1 enzymes. A) mRNA expression levels of Aldh1a1-3 enzymes relative to housekeeping genes in sorted stromal cells and dendritic cells of week 1 peripheral lymph nodes (pLN). B-D) Immunofluoresence analysis of Aldh1a1 expression at week 0 pLN (B), wk1 pLN (C) and wk6 pLN (D). The data represent n = 3. Error bars in A show mean ± SEM; * p <0.05, one-way ANOVA with Bonferroni’s multiple comparison test.

Since lymph node BECs are not recognized as Aldh1 expressing cells, we wanted to verify the expression by protein levels. We therefore performed immunofluorescence on post-natal developing lymph nodes and could show the expression of Aldh1a1 in BECs. Within BECs, Aldh1a1 expression could be detected already at day of birth (fig. 3B) and was abundantly present in BECs at day 7 after birth (fig. 3C). Upon analyzing Aldh1a1 enzyme expression over time, we observed weak Aldh1a1 expression in BECs in lymph nodes of week 6 old mice (fig. 3D). At that time, we observed Aldh1a1 expression on a subset of FDCs in B cell follicles. This expression was only seen in lymph nodes of adult mice (suppl. fig. 3).

Together, these data indicate that during early post-natal lymph node development, Aldh1 enzymes are expressed by multiple cell types that all could serve as cellular sources for retinoic acid that potentially mediate further FDC development.

### Postnatal abrogation of retinoic acid signaling in nestin-precursors does not prevent FDC formation

The multicellular distribution of retinoic acid producing enzymes imposes an impossibility to specifically assign the importance of these cellular sources for postnatal FDC development since the functional consequences of cell specific Aldh1 deletion in one cell subset, could be rescued by the expression of another Aldh1 enzyme in the same cell type (or by the expression of the same and other enzymes in another subset.

However, cell specific ablation of retinoic acid receptor signaling in precursor cells could be used to study the importance of retinoic acid signaling for further postnatal FDC development. Although the precursor for FDCs within neonatal lymph nodes have not been identified so far, we have shown in the past that nestin expressing cells can give rise to FDCs (Koning et al., 2016). We reasoned that ablation of retinoic acid receptor signaling in these cells shortly after birth would impact further FDC development. We performed cell specific ablation of retinoic acid receptor signaling in nestin expressing cells by crossing Nes-Cre^ERT2^ mice with RAR-DN mice in order to abrogate retinoic acid signaling specifically in nestin expressing cells upon tamoxifen treatment. Animals were treated from day 5 after birth for 5 consecutive days and lymph nodes were isolated when the animals were 2 or 3 weeks old.

Upon transcriptomic analysis of whole lymph nodes, we observed no difference in the mRNA expression of RARβ (data not shown) most likely because only a small subset of cells was targeted. mRNA expression of Cxcl13 however was reduced in peripheral lymph nodes of wk2 old animals (fig. 4A). This was mainly attributed to a reduction of Cxcl13 expression in axillary and brachial lymph nodes since similar levels of Cxcl13 were found in iLN of mice that were Nes-Cre^ERT2^ negative. (suppl. fig. 4A). Since CD35^pos^ FDCs were not present yet in both experimental groups (data not shown), we analysed the size of the B cell follicles, which showed no differences between lymph nodes from both groups (fig. 4B). In order to see whether FDCs would develop we examined mice that were left untreated for 3 weeks after 5 day tamoxifen treatment. We observed no differences in Cxcl13 mRNA expression (fig. 4C) and upon histological analysis FDCs could be found within B cell follicles of both groups (fig. 4D).

**Figure 4.**
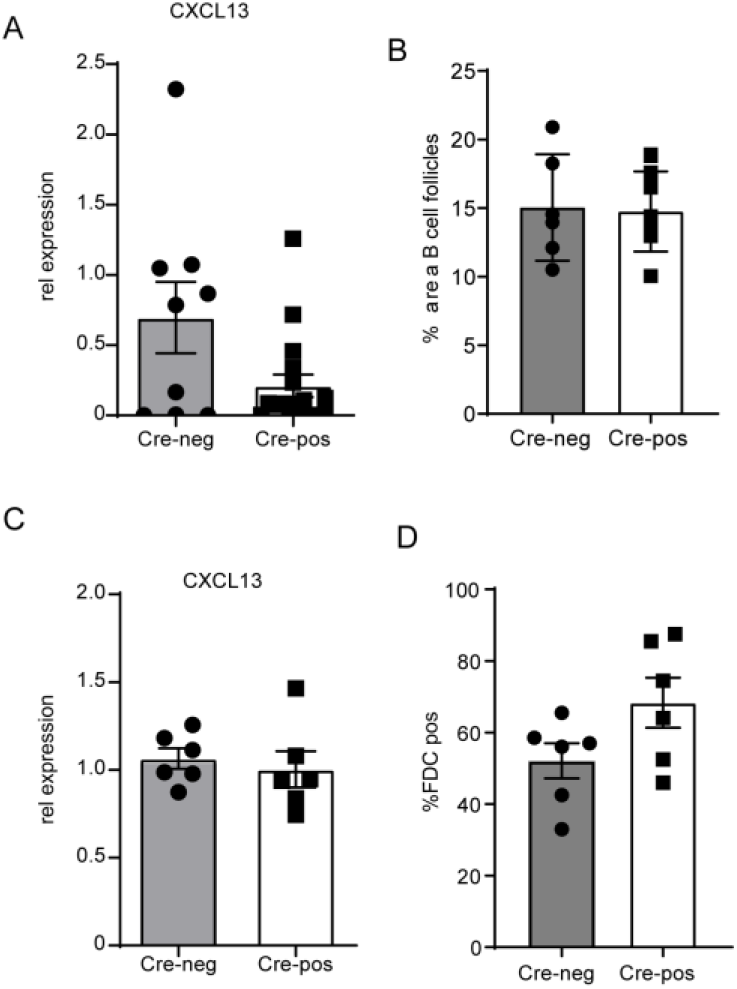
RAR signaling blockade in nestin precursors does not prevent FDC formation. A) Relative Cxcl13 mRNA expression in peripheral lymph nodes of 2 week old Nes-Cre^pos^ × RarDN mice (n=6) compared to Nes-Cre^neg^ × RarDN (n=3) littermates. B) Quantification of B cell areas in 2 week old Nes-Cre^pos^ × RarDN mice compared to Nes-Cre^neg^ × RarDN littermates. C) Relative Cxcl13 mRNA expression in peripheral lymph nodes of 3 week old Nes-Cre^pos^ × RarDN mice compared to Nes-Cre^neg^ × RarDN littermates (n=2). D) Quantification of %FDC pos B cell follicles in week 3 old Nes-Cre^pos^ × RarDN mice compared to Nes-Cre^neg^ × RarDN littermates (n=2). The data represent mean ± SEM. Dots or squares represent data of individual lymph nodes of the indicated number of mice.

Based on these observations, we can conclude that deleting retinoic acid signaling specifically in nestin precursor cells shortly after birth does not result in blockade of FDC development.

### Early postnatal blockade of RAR signaling prevents FDC development

In order to block all retinoic acid signaling and address whether retinoic acid mediated signaling is involved in FDC development postnatally, we performed pharmacological inhibition of retinoic acid receptor signaling using oral application of BMS493 (Mizee et al., 2013; S.A. van de Pavert et al., 2014). Hereto, we inhibited retinoic acid receptor signaling from postnatal day 4 onwards for 7 or 10 consecutive days.

As expected, transcript levels of RAR-β were down-regulated in peripheral as well as mesenteric lymph nodes upon BMS treatment at both timepoints indicating effective blockade of retinoic acid signaling (fig. 5A, suppl fig 5A). As a consequence, mRNA levels of Cxcl13 were also reduced and we hardly observed Cxcl13 protein expression in B cell areas and subcapsular sinuses of lymph nodes (fig. 5B and C). Importantly, other transcripts related to T cell zone stromal cells, such as Ccl19, Ccl21, Podoplanin (gp38), Cxcl12 and Il-7, were not significantly changed upon blockade of retinoic acid receptor signaling, indicating normal development of the other stromal cell subsets. As Mfge8 has been indicated as a marker for FDC precursors in the spleen (Krautler et al., 2012), we checked for its expression upon blockade of retinoic acid receptor signaling and observed that transcript levels were reduced upon BMS493 treatment (fig 5A).

**Figure 5.**
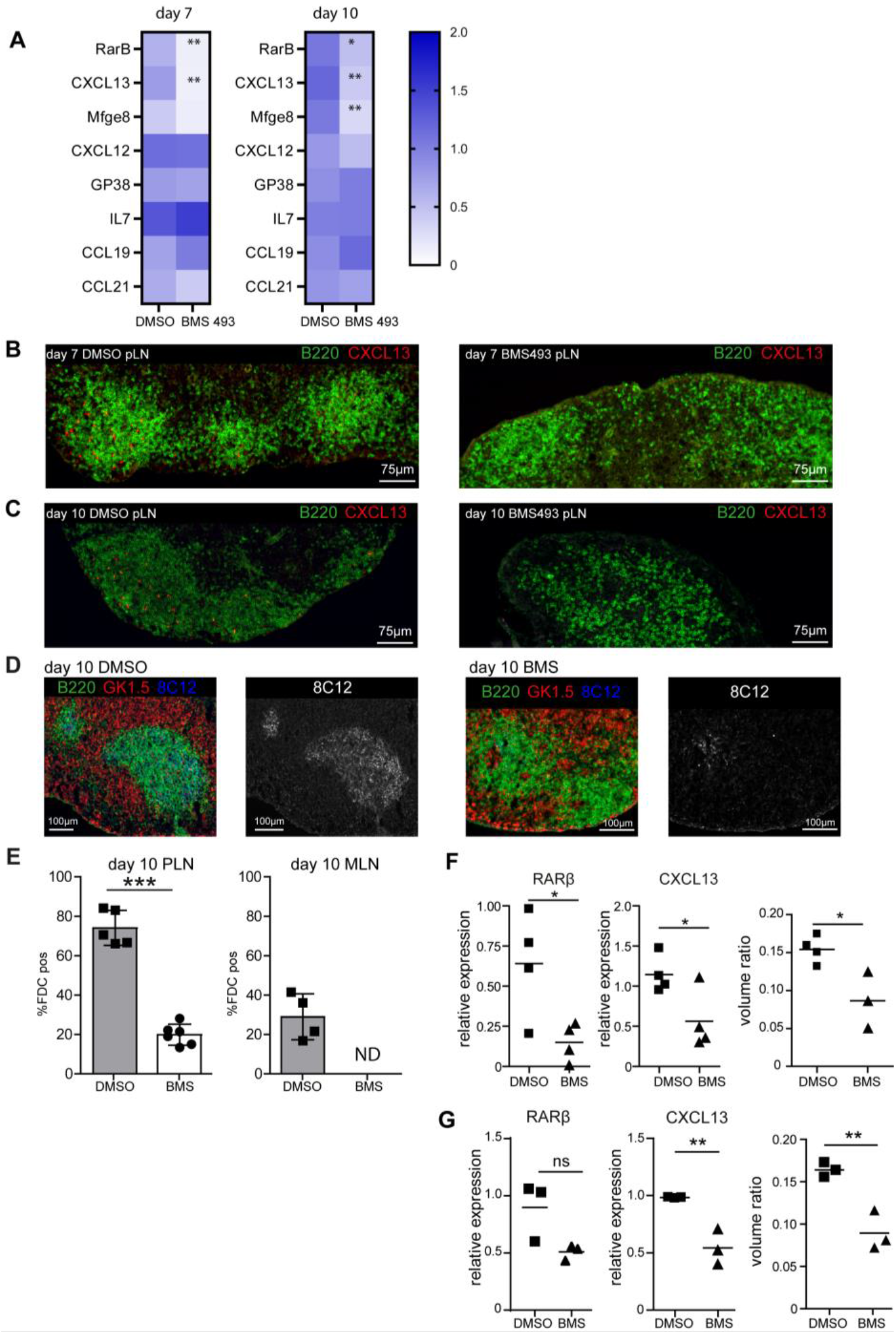
Inhibiting retinoic acid receptor signaling prevents CXCL13 expression and development of FDCs in lymph nodes. A) Heatmap showing mRNA expression levels of listed genes in peripheral lymph nodes upon treatment for 7 and 10 days with DMSO or BMS493, starting at day 4 after birth. B-C) Immunofluorescence analysis of CXCL13 protein expression (in red) in lymph nodes of DMSO (control) and BMS treated mice at day 7 (B) and day 10 (C) after start of treatment. D-E) Immunofluoresence analysis (D) and quantification (E) of FDCs (8C12 in blue) in lymph nodes of DMSO (control) and BMS treated animals at day 10 after treatment. F-G) mRNA expression levels of RARβ and CXCL13 and volume ratio of B cell follicle volume over total lymph node volume in peripheral lymph nodes of mice that were treated for 14 days with suboptimal doses of BMS493 vs DMSO (F) or for 7 days with BMS493 vs DMSO and left untreated for 21 days (G) starting at day 4 after birth. The data represent mean ± SEM; n = 3 or more, *, p < 0.05; **, p < 0.01; ***, p < 0.001; unpaired student’s *t* test. Squares and triangles in figures represent data of individual mice.

Most importantly, inhibiting retinoic acid receptor signaling through BMS493 treatment almost completely prevented the development of CD35^pos^ FDC networks in B cell areas in both peripheral and mesenteric lymph nodes (fig. 5D and E). In mesenteric lymph nodes, inhibiting RAR-signaling prevented the development of FDCs completely, while the percentage of B cell follicles that contained FDCs were significantly reduced in peripheral lymph node (fig. 5E).

The absence of Cxcl13 expression and FDC networks severely affected B-lymphocyte organization. In both peripheral and mesenteric lymph nodes of BMS treated mice, the number of B cell follicles that had formed at day 10 was lower compared to controls (suppl. fig. 5 C & D).

Within the spleen, transcripts for RAR-β were also reduced as a result of BMS493 treatment. At day 7 this resulted in a reduced transcript level of Cxcl13 while 10 days after treatment, Cxcl13 transcript levels were not different from control treated mice (suppl. fig. 5E). Immunofluorescence analysis revealed development of FDC networks in spleens from BMS treated mice (suppl fig. 5F), suggesting different signaling requirements for FDC development in spleen compared to lymph nodes.

To reduce, and not completely block, retinoic acid mediated signaling, we provided suboptimal doses of BMS493 starting at day 4 postnatal for 14 consecutive days. This treatment resulted in lower RAR-β and Cxcl13 transcript levels (fig. 5F) and we could detect low levels of Cxcl13 protein in the B cell follicles as well as FDC networks although they appeared smaller (data not shown). Whole lymph node imaging using light sheet microscopy, revealed that the total lymph node volume as well as the B cell follicle volume in suboptimal BMS493 treated peripheral lymph nodes were lower compared to control treated lymph nodes leading to a lower percentage the of LN volume that is occupied by B cell follicles (ratio of B cell area/total LN area) (fig. 5F and suppl fig. 5H-I). We observed similar results when we blocked FDC development for 7 days and left the mice untreated for 21 days (fig. 5G and suppl. fig. 5J).

In summary, these data show the requirement of retinoic acid receptor signaling for postnatal development of FDCs within lymph nodes.

## Discussion

Our results show the importance of retinoic acid signaling for the postnatal development of follicular dendritic cells (FDCs) and consequently B cell follicle formation in adult mouse lymph nodes.

Previous research has shown that lymphotoxin as well as TNFα mediated signaling is a prerequisite for the presence and maintenance of mature FDCs (Endres et al., 1999; Fu et al., 1998). Lymphotoxin signaling is not exclusively required for the differentiation and subsequent maintenance of FDCs as also FRCs and MRCs depend on signaling via this pathway (Roozendaal & Mebius, 2010). This suggests that additional signaling events are needed for precursors to develop into the various stromal subsets. Since retinoic acid plays a crucial role in the initiation of lymph node development before lymphotoxin mediated signaling takes place we hypothesized that retinoic acid signaling could be important for the early differentiation of mesenchymal precursors towards different stromal subsets as well.

Indeed, our results from *in vitro* stimulated mesenchymal-derived precursors showed that retinoic acid prevented the expression of transcripts that are normally induced upon LTβR signaling through activation of the non-canonical Nf-κB pathway, and that are associated with FRCs (Dejardin et al., 2002). It has been shown before that administration of retinoic acid leads to reduced Nf-κB activity *in vivo* without affecting p65 DNA binding capacity (Austenaa et al., 2004). We here show that retinoic acid selectively inhibits transcripts associated with T cell stroma, whose expression depends on the activation of the non-canonical Nf-κB pathway, without affecting relB or p52 DNA binding capacity. Even more so, Cxcl13 expression in mesenchymal precursors was unaltered upon sequential stimulation with retinoic acid and LTβR, suggesting a role for retinoic acid in favoring differentiation towards B cell stroma.

By blocking retinoic acid signaling *in vivo*, we were able to prevent the development of FDCs in lymph nodes, which has been shown to depend on lymphotoxin and TNFα mediated signaling. Together, our data points to a model in which mesenchymal precursor cells for FDCs are sequentially triggered, first by retinoic acid and subsequently by lymphotoxin and TNFα to finally differentiate towards FDCs.

This fate determining process potentially takes place before birth, since Cxcl13^high^Ccl19^neg/low^ cells could be observed at day of birth already indicating that differentially activated LNSCs are already present. Of interest is the observation that within developing B cell follicles, both Cxcl13 single positive, as well as Cxcl13/Ccl19 double positive cells were identified. Upon further development, these Cxcl13/Ccl19 double positive cells seems to disappear from the B cell follicles. Also after birth, retinoic acid signaling seems to be crucial for proper FDC development, since post-natal blockade with pharmacological inhibitors prevented development of FDC networks. Although we cannot prevent the appearance of Cxcl13^pos^ cells upon postnatal BMS493 treatment, due to their presence already at day of birth, their final maturation into mature FDCs was prevented. This suggests that after birth, retinoic acid signaling is a prerequisite for final FDC differentiation.

This neonatal fate determining step takes place when precursors are in close association with cells expressing Aldh1 enzymes which are dendritic cells, (lymphatic) endothelial cells as well as mesenchymal stromal cells.

In the absence of retinoic acid mediated signaling, precursors can still differentiate towards T cell stromal cells upon LTβR mediated signaling. Indeed, in the scenario in which retinoic acid mediated signaling was blocked, FDC and B cell follicles failed to develop while expression of FRC related chemokines was detected in normal amounts in stromal cells.

Within lymph nodes, expression of Aldh1 enzymes has been reported in gut derived DCs, lymph node stromal cells in mLN and epithelial cells in Peyer’s patches (Molenaar et al., 2011; Suzuki et al., 2010; Zhang et al., 2016). In addition, we identified endothelial cells, both high endothelial venules (HEVs) and lymphatic endothelial cells (LECs) as cells that transiently express Aldh1 enzymes most prominently during a period in which lymph nodes expand and stromal subsets are being formed. The observed decrease in Aldh1 expression after this period of expansion suggests that the differentiation of mesenchymal precursors towards the B cell stromal lineage occurs within a postnatal timeframe, which fades away in the absence of inflammatory stimulus once FDC networks are established. Whether endothelial cells re-express Aldh1 enzymes upon activation when FDC networks remodel and new FDCs are needed (Jarjour et al., 2014) has not been studied.

It is unknown how Aldh1 expression in neonatal lymph nodes is induced but (hematopoietic) cell subsets like the neo-migratory DCs (Zhang et al., 2016) could be involved. These cells express Aldh1 enzymes themselves and induce maturation of HEVs as their arrival in gut as well as skin draining lymph nodes leads to increased amounts of PNAd expression on HEVs. This migration depends on commensal fungi and absence of microbiota clearly affects their presence in peripheral lymph nodes (Zhang et al., 2016). Whether neo-migratory DCs are involved in inducing Aldh1 expression on HEVs and LECs and whether retinoic acid produced by these DCs are in fact instrumental for this effect is unknown. However, the blockade of retinoic acid signaling that prevented FDC development in the experiments presented here occurred before the entry of neo-migratory DCs in skin draining lymph nodes, which is around 2 weeks after birth, suggesting that they are not involved (Zhang et al., 2016).

While the first description of FDCs has been a long time ago already (Mitchell & Abbot, 1965), the discovery of their precursors in lymphoid organs has remained elusive until recently. Whereas in the spleen specific precursors for FDCs have been documented in several reports (Castagnaro et al., 2013; Krautler et al., 2012), progenitor-progeny relationships for lymph nodes are not so clear yet. Lineage tracing studies showed MRCs at least in part to be precursors for FDCs (Jarjour et al., 2014). Other models have shown that upon neonatal lineage tracing of lymphoid tissue organizer cells, a subset of these cells give rise to MRCs, but not to FDCs

(Hoorweg et al., 2015). In addition, as mentioned by Jarjour et al. other precursors for FDCs may exist (Jarjour et al., 2014). In the spleen, precursors for FDCs can be identified based on the expression of Mfge8 and Cxcl13 and these cells are perivascular located. Here we show that also in neonatal lymph nodes, Mfge8^pos^Cxcl13^pos^ cells can already be identified at day of birth throughout the lymph node and over time, these cells start to localize in B cell follicles as well as in the subcapsular sinus. In adult lymph nodes, the majority of these cells can be detected by the expression of CD35, a marker for FDCs indicating that lymph node FDC precursors can be identified by similar markers as well. However, in adult lymph nodes, we did not detect Mfge8^pos^CXCL13^pos^ expressing cells surrounding blood vessels. This could mean that either precursors for FDCs are not located perivascular or that they do not express these markers upon homeostasis.

Our results allow for a lineage tree model in which precursors upon encounter with retinoic acid and subsequent stimulation with lymphotoxin will become a functional FDC, while they differentiate towards FRC in the absence of retinoic acid mediated signaling.

## Materials and methods

### Mice

C57BL/6 were bred and maintained at the Amsterdam Animal Research Center (AARC) under specific pathogen free conditions. The Animal Experiments Committee of the VU (Vrije Universiteit) University Medical Center approved all of the experiments described in this study. RAR DN403 (a kind gift of Dr. S. Sockanathan, Dept of Neuroscience, Johns Hopkins School of Medicine, Baltimore (Rajaii, Bitzer, Xu, & Sockanathan, 2008)), and Nes-cre^ERT2^ (C57BL/6-Tg(Nes-cre/ERT2)4Imayo (Imayoshi, Ohtsuka, Metzger, Chambon, & Kageyama, 2006)) mice were bred and maintained at Palo Alto Veterans Institute of Research, VA Palo Alto Health Care System, Palo Alto, CA 94304, USA.

### BMS treatment

For *in vivo* blockade of retinoic acid signaling, mice were treated with the pan-RAR antagonist BMS493 (5mg/kg; Tocris Bioscience) or vehicle (DMSO) in corn oil as described before (S.A. van de Pavert et al., 2014). Mice were treated for 7 or 10 days via oral gavage twice daily with 10-12 hrs intervals starting at day 4 after birth. After 7 or 10 days, mice were sacrificed for analysis. For temporal blockade of retinoic acid signaling mice were treated the same way for 7 days and left untreated for another 21 days after which they were sacrificed for analysis. For suboptimal blockade of retinoic acid signaling, mice received 2,5mg/kg BMS493 or vehicle (DMSO) for 14 days. Per experimental group (BMS493 vs vehicle) per time point, all animals within 1 litter either receive BMS493 or DMSO, to prevent contamination of of the treatment within one litter. Litters were randomly allocated in experimental groups.

### Tamoxifen treatment of Nes-cre^ERT2^ × RAR DN403 mice

Tamoxifen (Sigma) was dissolved in ethanol first and subsequently preheated corn oil (42°C, Sigma) was added and strongly mixed for 1 hr in the darkness. Final concentration was 10mg/kg. For postnatal induction, tamoxifen solution was applied *i.p.* to the mothers (100 μl/ day) starting at p4 (Schultheiss et al., 2013). Animals were sacrificed at day 14 (p14) or day 21 (p21).

### Single cell suspensions

E13.5 mesenchymal cells were prepared as described (S. A. van de Pavert et al., 2009). In short, the head, extremities and organs were removed from E13.5 embryos and the remainings were enzymatically digested with Blendzyme 2 (0,5 mg/ml; Roche Applied Sciences, Almere, The Netherlands), DNA-se I (0,2 mg/ml; Roche) in PBS for 15 min at 37°C while continuous stirring. Cell suspensions were washed with RPMI supplemented with FCS (2%) as well as antibiotics and counted. Subsequently, cells were cultured in Mesencult proliferation medium (Mesencult Basal Medium supplemented with MSC stimulatory supplements (StemCell Technologies, Grenoble, France) and 2% antibiotics and glutamine). Upon their second passage, cells were used for experiments.

Single cell suspensions of lymph nodes were generated as described (Fletcher et al., 2011). In short, upon isolation, lymph nodes were pierced with a 25g needle and place in ice-cold RPMI1640 (Invitrogen). LNs were subsequently digested using an enzyme mixture of 0.2 mg/ml collagenase P (Roche), 0.8 mg/ml Dispase II (Roche) and 0.1 mg/ml DNase I (Roche) in RPMI medium (Invitrogen) without serum. With 5 min intervals, contents were gently mixed. After 15 min, suspensions were gently resuspended, supernatant collected in ice-cold PBS (2% FCS and 5mM EDTA) and centrifuged (5 min 300g, 4°C). Fresh enzyme mix was added to the tubes and digestion was repeated as described. After 15 min, supernatant was collected in the same tube with ice-cold PBS and fresh enzyme mix was added to the remaining fragments. Every 5 min, suspensions were mixed vigorously till no fragments remained. Supernatant was collected in the same collection tube and centrifuged (5min, 300g, 4°C).

### Differentiation assays

For adipogenesis, 2×10^5^ cells/well in a 6-well plate were incubated with DMEM supplemented with 10% FCS, 1% antibiotics, 1μM dexamethason (Sigma, Sigma-Aldrich Chemie B.V, Zwijndrecht, The Netherlands), 0.1mM indomethacin, 0.5 mM 3-isobutyl-1methylzanthine (IBMX, Sigma) and 10μM insulin (Sigma). Medium was replaced 2 times per week for 3 weeks, after which the cells were fixed with 10% formaldehyde for 20 minutes at RT and stained with Oil-red-O (Sigma). For chondrogenesis, 4×10^5^ cells were pelleted in a 15 ml tube and cultured in DMEM supplemented with 1% antibiotics, ITS-premix (BD Biosciences, Becton Dickinson B.V., Breda, The Netherlands), 0,1 μM dexamethasone, 86.5μM L-ascorbic acid 2-phosphate (Sigma) and 10ng/ml TGF-ß1 (Peprotech, London, UK). For microscopy, pellets were sectioned and stained for 10 minutes with 0.2% Toluidine Blue. For osteogenesis, 50×10^3^ cells/well were cultured in a 6-well plate with DMEM supplemented with 10% FCS, 1% antibiotics, 0.1μM dexamethasone, 10mM ß-glycerophosphate (Sigma) and 173μM L-ascorbic acid 2-phosphate. Cells were fixed for 15 minutes in 10% formaldehyde and subsequently stained with Alizarin-Red-S (20mg/ml, BDH, United Kingdom).

### In vitro stimulation and mRNA transcript analysis

To determine transcript levels, 1×10^5^ mesenchymal cells were seeded in 24-wells plate and allowed to adhere for 2-4 hrs in 1:1 (vol/vol) mesencult proliferation medium and DMEM-F12 medium supplemented with 10 % FCS and 2 % antibiotics with glutamine. After 2-4 hrs, medium was replaced with DMEM-F12, 5% FCS with 2% antibiotics and glutamine. The next day, stimulations with retinoic acid or agonistic anti-LTβR mAb were performed in duplo in DMEM-F12, 2% FCS with 2% antibiotics and glutamine. For sequential stimulation experiments, cells were cultured for 6 hrs either in medium alone or in the presence of retinoic acid (100 nM, Fluka, Sigma-Aldrich, Zwijndrecht, The Netherlands, dissolved in ethanol), after which the cells were washed and stimulated with or without agonistic anti-LTβR mAb for another 6 hrs (2μg/ml, clone 4H8-WH2, produced in Carl Ware’s laboratory) in duplo. Cells were subsequently lysed, after which mRNA was isolated from total RNA using the mRNA capture kit (Roche) and cDNA was synthesized using the Reverse transcriptase kit (Promega Benelux, Leiden, The Netherlands) according to manufacturer’s protocol.

To determine transcript levels within whole mount lymph nodes, lymph nodes were harvested, lysed and homogenized in TRIzol Reagent (Life Technologies). RNA was isolated by precipitation with isopropanol according to the manufacturer’s protocol and cDNA was synthesized from total RNA using RevertAid First Strand cDNA Synthesis Kit (Fermentas Life Sciences) according to the manufacturer’s protocol.

Quantitative RT-PCR was performed in duplo on an ABI Prism 7900HT Sequence Detection System (PE Applied Biosystems) or StepOnePlus Real-Time PCR System (ThermoFisher Scientific). Total volume of the reaction mixture was 10 μl, containing cDNA, 300nM of each primer and SYBR Green Mastermix (PE Applied Biosystems). From a set of 8 housekeeping genes, the two most stable were selected (Cyclo and HPRT). The comparative Ct method (∆Ct) was used to indicate relative changes in mRNA levels between samples. Relative mRNA levels of unstimulated cells or control treated tissues were set at 1,0.

### Flow cytometric analysis and cell sorting

To determine cell surface expression, 1×10^5^ cells were cultured as indicated. Cells were stimulated with retinoic acid (100 nM), the agonistic anti-LTβR mAb or both for 24 hrs and 72 hrs. Cells were harvested by trypsinization and subsequently stained with biotin conjugated anti-VCAM-1 or anti-ICAM-1 (both eBioscience, Immunosource, Halle-Zoersel, Belgium), unlabeled anti-MAdCAM-1 (clone MECA 367, affinity purified from hybridoma cell culture supernatants) and unlabeled anti-LTβR mAb and subsequently counterstained with streptavidin-Alexa 488 and goat-anti-rat-Alexa 488 (all Invitrogen, Breda, The Netherlands), respectively. Sytox Blue (Invitrogen) staining was used to discriminate between live and dead cells. Data were acquired on a Cyan ADP High Performance Research Flow Cytometer (Beckman Coulter) and were analyzed with Summit Software v4.3.

For cell sorting, single cell suspensions of lymph nodes were stained in 100 μl diluted antibody mixtures containing the following antibodies; CD45-PE-Cy7 (Biolegend), Ter119-BV605 (Biolegend), GP38-Alexa Fluor 488 (Biolegend), CD31-PE (Biolegend), MHC-II-647(Clone M5-114, MO2Ab facility) and Cd11c Apc-Cy7 (Biolegend). Cells were sorted on a BD Fusion (BD Biosciences) using a 100 μm nozzle. Sorted cells were collected in ice-cold HBSS (Invitrogen), centrifuged and lysed in Trizol.

### RelB translocation and TransAM-analysis

Nuclear translocation of relB was determined on cytospins of cells stimulated for 6 hrs with retinoic acid followed by agonistic anti-LTβR mAb stimulation for 6 hrs. Cells were fixed with 4% paraformaldehyde for 5 min and subsequently permeabilized with 0.2% Triton-X and stained with anti-relB (Cell Signaling Technology, Danvers, USA) followed by goat-anti-rabbit Alexa 546 (Invitrogen). Cells were embedded in Vinol mounting media (Air Products, Allentown, USA) supplemented with DAPI (Invitrogen) to visualize nuclei. Analysis was performed on a Leica DM6000 fluorescence microscope (Leica Microsystems) equipped with LAS AF software.

For TransAM analysis, cells were stimulated for 6hrs with retinoic acid (100 nM), followed by 3hrs stimulation with the agonistic anti-LTβR mAb, after which nuclear extracts were prepared with the Nucbuster protein extraction kit (Thermo Scientific, Rockford, USA). DNA binding capacity of the Nf-κB subunits relB and p52 were determined with the Nf-κB TransAM family kit (Active Motif, Rixensart, Belgium) according to manufacturer’s protocol.

### Fluorescent in situ hybridization

Lymph nodes were fixed for 2 hrs in 4% paraformaldehyde at RT. Samples were cryoprotected and subsequently embedded in OCT compound (Sakura Finetek Europe) and stored at −80°C until sectioning all under RNAse free conditions. Cryostat sections (7μm) were collected on superfrost plus adhesive slides (VWR, the Netherlands) and multiplex *in situ* hybridizations were performed according to manufacturer’s protocol (Molecular Instruments and described in (Choi et al., 2018)). In short, slides were washed with 5x SSCT, pre hybridized in hybridization buffer at 37°C in a humidified chamber. Probes were added at 0.4 pmol/100 μl in probe hybridization buffer at 37°C and hybridized overnight in a humidified chamber, while covered with parafilm. Next day, slides were washed in decreasing concentrations of probe wash buffer in 5x SSCT (100%, 75%, 50%, 25% respectively) and washed in 5x SSCT at room temperature. Next, samples were pre-incubated with amplification buffer for 30 min. Hairpin solutions were prepared with 6 pmol snap-cooled hairpins/100 μl and added to the slides. Samples were incubated overnight at room temperature in a humidified chamber. Next day, excess hairpins were removed by washing with 5x SSCT. In case of additional antibody staining, antibodies and/or nuclear labeling were added during these washing steps. Finally, slides were embedded in mounting medium and stored at 4°C till analysis.

### Immunofluorescence

E18.5 embryos were fixed in 0.4% formaldehyde overnight at 4°C, lymph nodes (postnatal day 0 – 6 weeks of age) and spleens from C57BL/6 J and Nes-CRE^ERT2^ × DN RAR 403 mice were fixed in 4% formaldehyde for 10 min. Samples were cryoprotected and subsequently embedded in OCT compound (Sakura Finetek Europe) and stored at −80°C until sectioning. Cryostat sections (7μm) collected on gelatin coated glasses were fixed in ice-cold acetone for 10 minutes and blocked with 10% NMS in PBS prior to antibody staining. Immunofluorescence staining was performed in PBS, supplemented with 0.1% (wt/vol) Bovine Serum Albumin (BSA).

Sections were embedded in Vinol + DAPI and analyzed on a Leica SP8Confocal Laser Scanning Microscope or a Leica DM6000 (both Leica Microsystems Nederland b.v., Rijswijk, The Netherlands).

For whole mount immunofluorescence, lymph nodes were fixed in 4% formaldehyde for 10min, washed with PBS, dehydrated with methanol series (50%, 75%, 95%, 100% (2x)) and subsequently rehydrated. Lymph nodes were blocked with PBS-MT (1% skim-milk, 0.4% TritonX-100) o/n at 4°C and subsequently incubated with directly labeled primary antibody for 4 days in PBS-MT, 4°C, rolling. After 3 washing steps in PBS, lymph nodes were embedded in 1,5% low-melting agarose for easy handling. Samples were dehydrated with methanol series (50%, 75%, 95%, 100%, 100%) and overnight incubated in 1:1 methanol : BABB (BenzylAlcohol-BenzylBenzoate 1:2, both Sigma)). Next morning, all solutions were replaced with BABB and stored in dark until acquisition. Acquisition was performed using the Ultramicroscope (La Vision BioTec, Bielefeld). Images were analyzed using Imaris Software (Bitplane, version 8.02 or higher).

For volumetric analysis of B cell follicles and lymph node size, individual B cell follicles as well as the whole lymph node were masked in Imaris Software and volumes were extracted using Imaris Vantage module. The ratio of B cell follicles within total lymph nodes was calculated as the total volume of B cell follicles within the lymph node volume.

The following anti-mouse antibodies were used; unlabeled anti-MECA325, anti-Aldh1a1 (Abcam) (van de Pavert et al., 2014), biotinylated anti-CXCL13 (R&D Systems), alexa fluor 488 labeled anti-B220 (clone 6B2), anti-MAdCAM-1 (clone MECA367), anti-CD4 (clone GK1.5), alexa fluor 555 labeled anti-CD4 (clone GK1.5), BV510 and alexa fluor 647 labeled anti-CD35 (clone 8C12, BD Biosciences), anti-CD31 (clone ERMP12) and eFluor 660 labeled anti-Lyve1 (clone Aly7, eBioscience).

Unconjugated antibodies were detected with species specific secondary reagents. Biotinylated anti-CXCL13 was visualized with signal amplification using a TSA^™^ Kit with HRP-streptavidin and Alexa Fluor 546 tyramide (Invitrogen).

### Statistical Analysis

Statistical analysis was performed as described in the figure legends. The letter ‘n’ in the figure legends, refers to the number of individual samples, or independent *in vitro* experiments. For all BMS493 animal experiments, sample size was estimated using a power analysis to verify that the sample size gave a value of > 0.9 if P was > 0.05. For all other experiments, no statistical method was used to predetermine sample size, experiment blinding upon analysis was not used.

---

**Supplementary figure 1.**
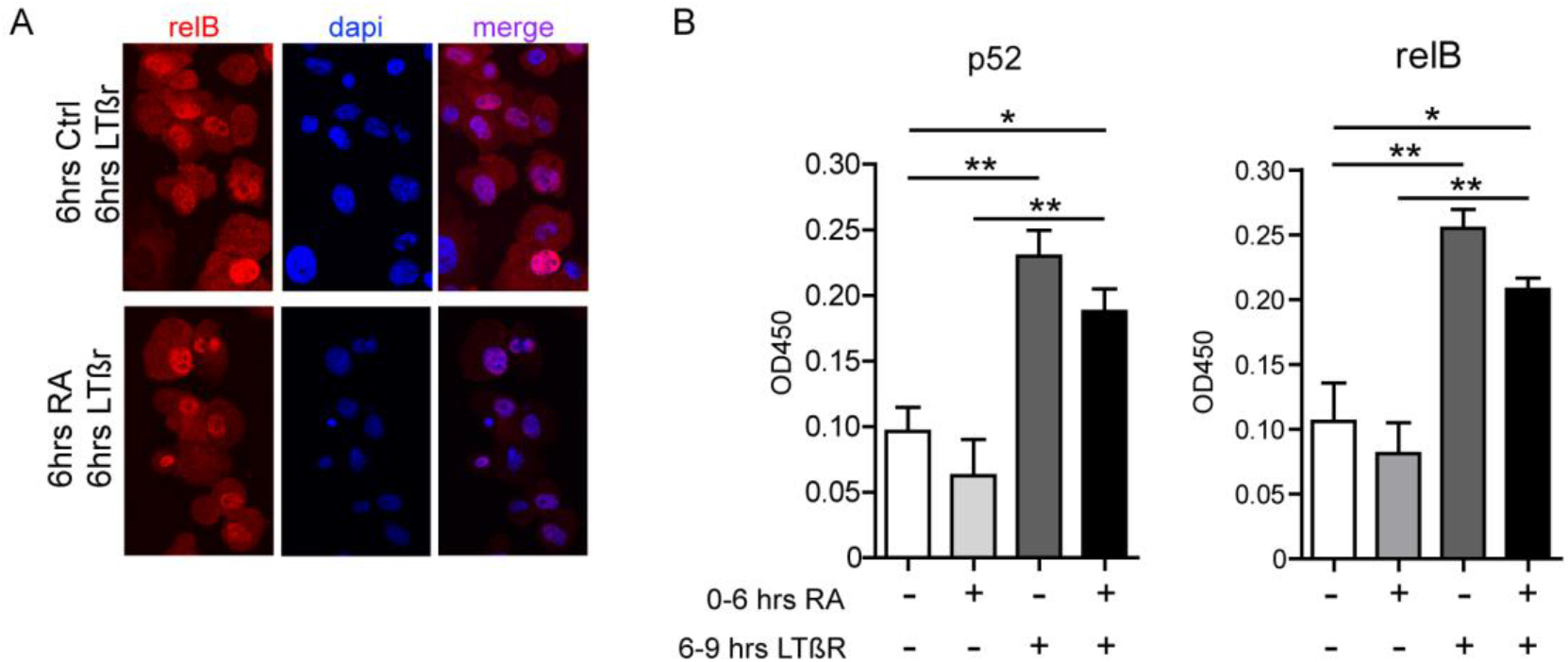
Related to Figure 1. *Retinoic Acid signaling prevents upregulation of FRC related genes*. A) Immunofluorescence staining for RelB (in red) and nuclei (in blue) in cytospins of mesenchymal precursors stimulated with or without retinoic acid (6hrs) followed by agonistic anti-LTβR mAb (6hrs). B) TransAM analysis of nuclear p52 and relB after stimulation of mesenchymal precursors with retinoic acid (6hrs) followed by agonistic anti-LTβR mAb (3hrs) or retinoic acid (6hrs) alone or anti-LTβR mAb (3hrs) alone. Results are representative of 3 independent experiments. The data represent mean ± SEM; n = 3, * p <0.05, ** p <0.01, one-way ANOVA with Bonferroni’s multiple comparison test.

**Supplementary figure2.**
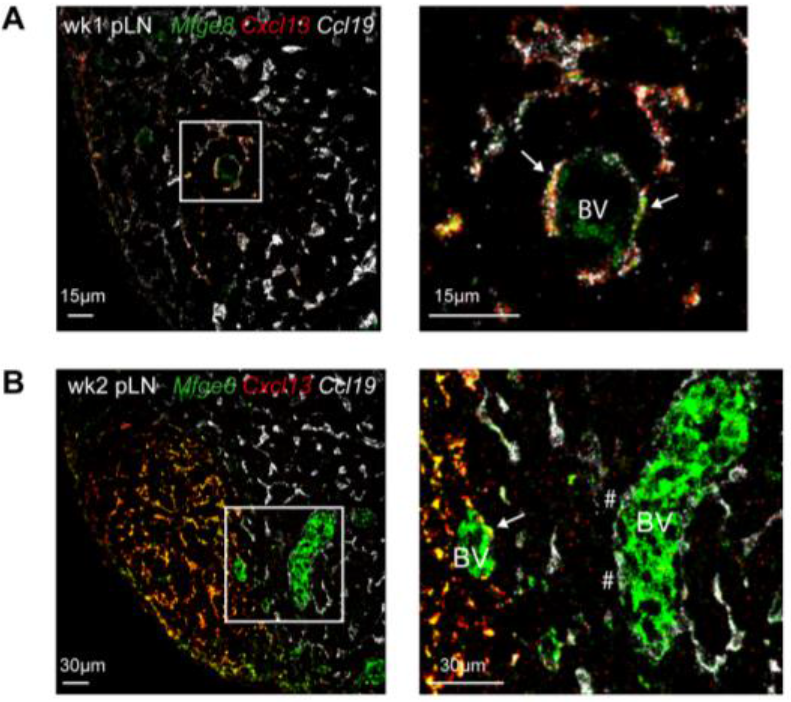
Related to Figure 2 *Localization of FDC precursors*. A-B) Confocal analysis of peripheral lymph nodes (pLN).by multiple fluorescence *in situ* hybridizations at different timepoints after birth A) Combined expression of Mfge8, Cxcl13 and Ccl19 in peripheral lymph nodes at 1 week after birth. Arrows indicate cells around blood vessel (BV) expressing Mfge8, Cxcl13 and Ccl19.B) Combined expression of Mfge8, Cxcl13 and Ccl19 in lymph nodes at 2 weeks after birth. Arrows indicate cells around blood vessel (BV) that express Mfge8 and Cxcl13, hashtags indicate cells around blood vessel (BV) that express Mfge8 and Ccl19. The data represent n = 3.

**Supplementary figure3.**
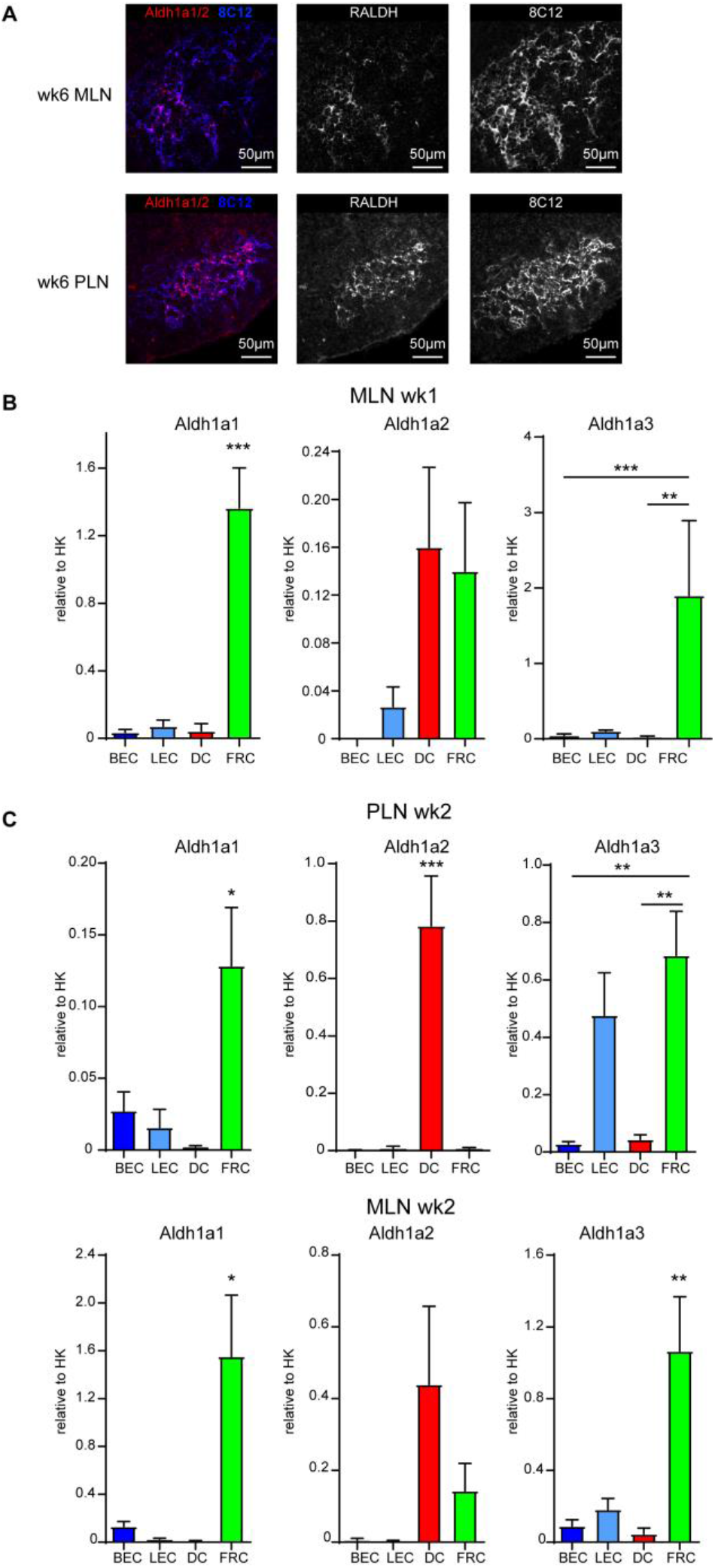
Related to figure 3 *Multiple cellular sources of Aldh1 enzymes*. mRNA expression levels of Aldh1a1-3 enzymes relative to housekeeping genes in sorted stromal cells and dendritic cells of peripheral lymph nodes (pLN) or mesenteric lymph nodes (mLN) at the indicated timepoints. The data represent mean ± SEM; * p <0.05, ** p <0.01, *** p <0.001 one-way ANOVA with Bonferroni’s multiple comparison test. n = 3 or 4.

**Supplementary figure4.**
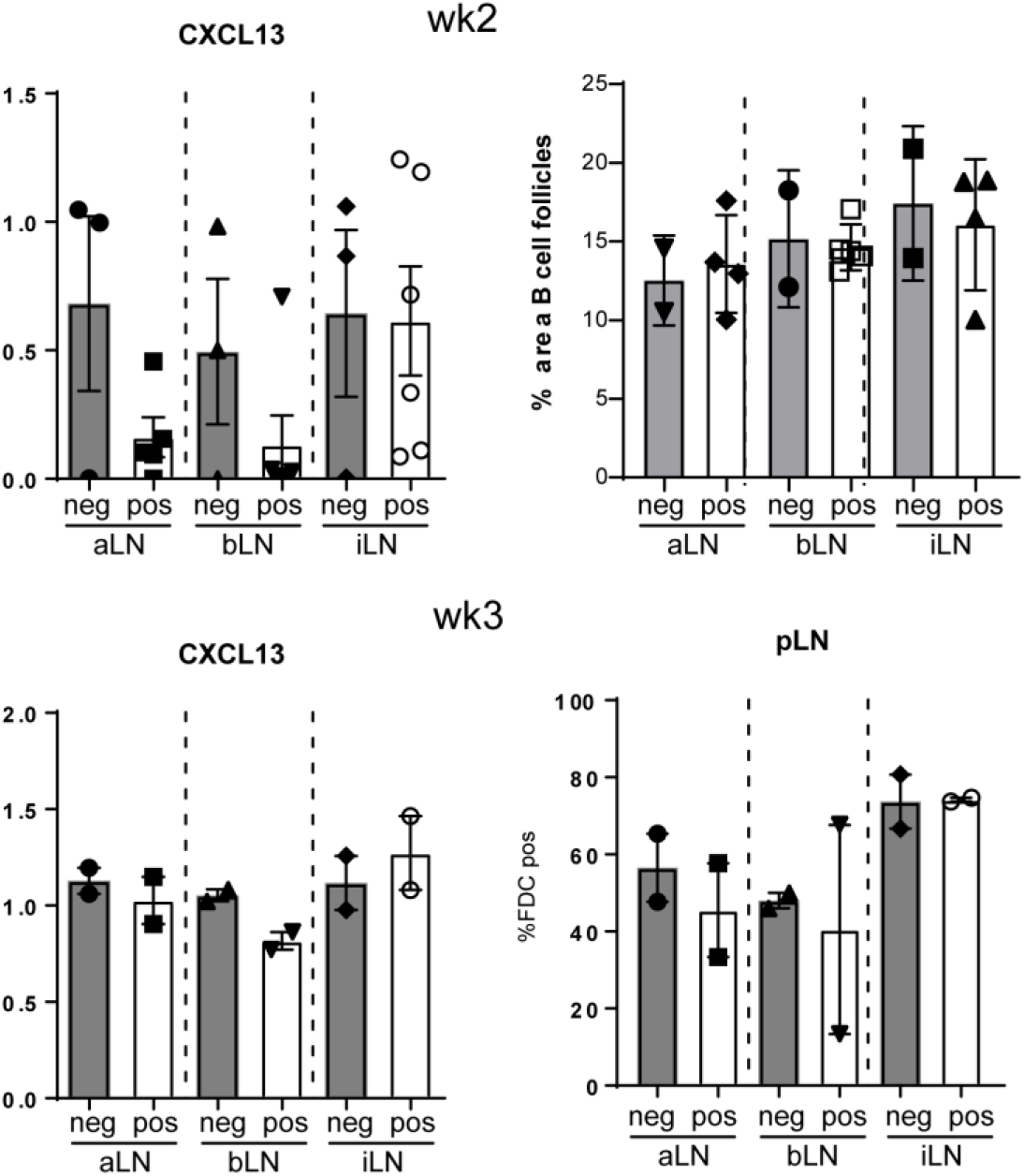
Related to figure 4. *RAR signaling blockade in nestin precursors does not prevent FDC formation.* A and C) Relative Cxcl13 mRNA expression in indicated peripheral lymph nodes (pLN) of Nes-Cre^ERT2pos^ × DN-RAR mice compared to Nes-Cre^ERT2neg^ × DN-RAR littermates at 2 weeks (A) or 3 weeks (B) after birth. B) Quantification of B-cell areas in 2 week old Nes-Cre^ERT2pos^ × DN-RAR mice compared to Nes-Cre^ERT2neg^ × DN-RAR littermates. D) Quantification of %FDC pos B-cell follicles in week 3 old Nes-Cre^ERT2pos^ × DN-RAR mice compared to Nes-Cre^ERT2neg^ × DN-RAR littermates. The data represent mean ± SEM; n = 2 or more. Dots or squares represent data of individual lymph nodes.

**Supplementary figure5.**
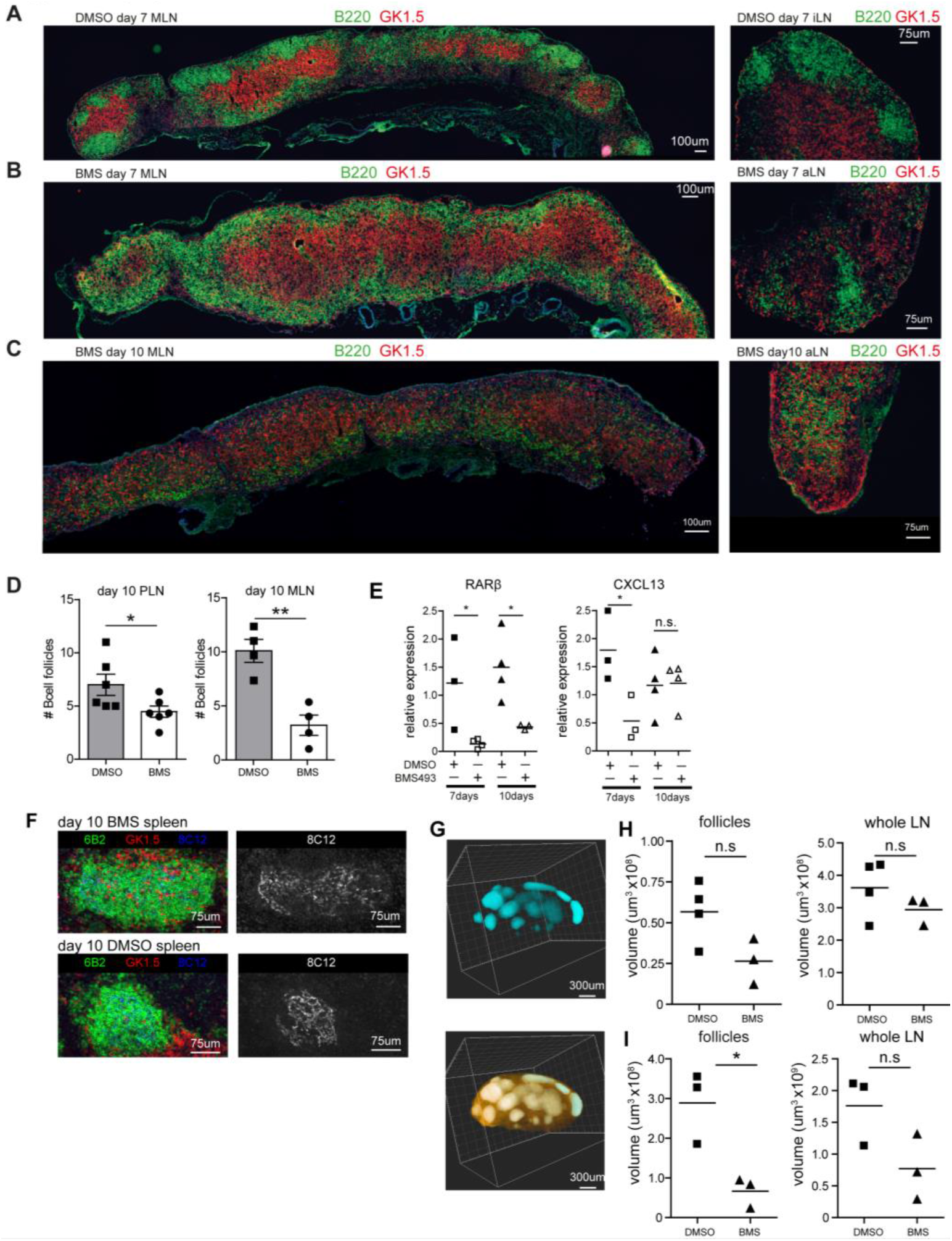
Related to Figure 5. *Endothelial cells are the source of retinoic acid during lymph node development*. A-C) Immunofluorescence staining for B cells (green) and T cells (red) in peripheral and mesenteric lymph nodes of vehicle treated (DMSO) and BMS493 treated animals. D) Quantification of the number of B cell follicles in DMSO and BMS493 treated animals in both peripheral and mesenteric lymph nodes. E) mRNA expression levels of RARβ and CXCL13 in spleen upon treatment with vehicle (DMSO) or BMS493 for 7 and 10 days starting at day 4 after birth. F) Immunofluorescence analysis of FDC (8C12 in blue) network in spleen of vehicle (DMSO) or BMS493 treated animals 10 days after treatment. G) Representative ultramicroscopy image of a whole mount stained peripheral lymph node from DMSO treated mice showing B cell follicles in blue and total lymph node volume in orange. H-I) Volume of B cell follicles and whole lymph nodes of peripheral lymph nodes of mice that were treated with suboptimal dose of BMS493 vs DMSO for 14 days starting at day 4 after birth (H) or for 7 days with BMS493 vs DMSO and left untreated for 21 days (I) Results are representative of at least 3 individual mice. The data in D,E,H-I represent mean ± SEM; n = 3. n.s. not significant. *, p < 0.05; ** p <0.01, unpaired student’s *t* test. Squares and triangles in figures represent data of individual mice.

